# Integrated Multi-omics Prioritizes Ribosomal-targeting Omacetaxine Mepesuccinate for Optic Neuritis Therapy

**DOI:** 10.1101/2025.07.22.666225

**Authors:** Yan Zhang, Maosheng Guo, Yihong Huang

## Abstract

**Background:** Optic neuritis (ON), an acute demyelinating disorder often preceding multiple sclerosis, lacks therapies preventing neurodegeneration despite corticosteroid use.

**Methods:** We combined GWAS (FinnGen cohort, finn-b-H7_OPTNEURITIS), immune-specific eQTL analysis (CAGE.sparse microglia/astrocytes), and network pharmacology. SMR-HEIDI filtering (p_HEIDI>0.05; FDR<0.05) validated causal genes.

**Results:** 46 ON-associated genes identified, with HLA-DRB1 as top risk locus (OR=3.297, FDR<0.01).Functional enrichment revealed antigen presentation dysregulation (GO:0002483, FDR=0.0025) and phagosome activation (hsa04145, FDR=0.0468), confirming microglial-astrocytic pathology.Omacetaxine mepesuccinate prioritized as sole repurposing candidate (FDR=0.0467) targeting ribosomal proteins RPL3/RPL2—mechanistically linked to glial protein synthesis suppression in demyelination.

**Conclusions:** This first multi-omics analysis of ON bridges HLA-mediated autoimmunity with repurposed ribosomal-targeted therapy, proposing omacetaxine for neuroprotection.

## Introduction

Optic neuritis (ON), an acute inflammatory demyelinating disorder of the optic nerve, represents the initial manifestation of multiple sclerosis (MS) in 25-50% of cases[1,2].Its annual incidence exhibits geographic variability (e.g., 1.2-7.5/100,000 in European populations) [1,3], with acute unilateral vision loss and periocular pain severely impacting patients’ functional capacity[4].

Current first-line therapy employs high-dose intravenous corticosteroids[1], which accelerate short-term visual recovery but fail to prevent retinal neurodegeneration[5].Optical coherence tomography (OCT) studies demonstrate persistent ganglion cell layer thinning (>20% reduction vs. baseline) after steroid treatment[6].Alternative interventions (e.g., plasma exchange, intravenous immunoglobulin) exhibit limited efficacy and lack Phase III evidence[3,7], underscoring an urgent need for mechanism-driven therapies targeting ON-specific pathways.

Genetic susceptibility plays a pivotal role in ON pathogenesis, with HLA-DRB1 alleles implicated in both ON onset and progression to MS [8,9] and HLA-DRB115:01, conferring 3.9-fold increased ON risk[10].Recent advances in immunogenomics have begun to unravel the molecular interplay between antigen presentation dysregulation and autoimmune demyelination [11][12],yet no existing therapies target these mechanisms.

To bridge this translational gap, we integrated three methodologies:Genome-wide association study (GWAS) of Finnish cohorts (FinnGen R10, finn-b-H7_OPTNEURITIS);Cell-type-specific expression quantitative trait loci (eQTL) analysis using microglia/astrocyte transcriptomes (CAGE.sparse);Network pharmacology screening for FDA-approved agents targeting causal pathways.This framework aims to identify repurposable neuroprotective therapies for ON, addressing an urgent unmet need.

## Materials and Methods

### 1. Genetic Data Acquisition

GWAS summary statistics for optic neuritis (FinnGen ID: finn-b-H7_OPTNEURITIS) were retrieved from IEU OpenGWAS (gwas.mrcieu.ac.uk). This dataset comprises 582 acute/subacute ON cases and 217,491controls from the Finnish biobank (Release R10), reflecting genotype-clinical diagnosis correlations in Finnish cohorts. Acute/subacute ON cases were prioritized to align with diagnostic consensus[10].

### 2. SMR-HEIDI Analysis

Summary-data-based Mendelian randomization (SMR) was performed using eqtlSMR v0.98 to infer causal genes, with parameters validated in neuroinflammatory disorders:

#### eQTL source

CAGE.sparse (population-based eQTL data from immune-relevant tissues, including astrocytes, microglia, and oligodendrocytes—key cell types in ON neuroinflammation[9].

#### Significance thresholds

p_SMR < 0.05(SMR test significance);p_HEIDI > 0.05(ensuring colocalization and excluding pleiotropy);nsnp_HEIDI ≥ 3(minimum SNPs for robust HEIDI testing);FDR < 0.05(Benjamini-Hochberg correction for multiple testing).

#### Visualization Procedure

Volcano plot generation was performed using R v4.2.1 (R Core Team, 2022) with the ggplot2 package v3.4.4 (Wickham, 2016). Statistical Significance: Adjusted p-value < 0.05 (Benjamini-Hochberg FDR correction applied).log_2_FC ≥ 1 are up-regulated genes,log_2_FC ≤ −1 are down-regulated genes.

### 3. Functional Enrichment Analysis

Genes surpassing SMR thresholds underwent functional annotation via: Gene Ontology (GO) was including Biological Process (BP), Cellular Component (CC), and Molecular Function (MF) terms;and,KEGG pathway analysis.Both analyses used the R package clusterProfiler (v4.6.0).Significantly enriched terms (FDR < 0.05) were visualized using ggplot2 (v3.4.4).

### 4. Network Pharmacology Drug repurposing leveraged

#### Drug database

DrugBank v5.1.9 (annotated with >14,000 targets). **Algorithm:** Proximity-based drug-disease association scoring via the network method published by Guney et al., Nat. Commun in 2016[12].Network topology analysis and visualization used igraph (v1.5.0) in R.

#### Statistical filtering

FDR<0.05 for significant drug candidates.

### 5. Ethics Declaration

All data were sourced from publicly available repositories:Genetic summary statistics: FinnGen R10 (IEU OpenGWAS ID: finn-b-H7_OPTNEURITIS, accessed 10/04/2025);Drug-target interactions: DrugBank v5.1.9 (accessed 26/04/2025) .No individual-level patient data were accessed or used in this study.

## Result

### 1 Identification of Optic Neuritis-Associated Genes

SMR-HEIDI analysis initially revealed 46 probe-gene associations (Supplementary Table 1). After probe deduplication—where for genes with multiple probes, the probe with the smallest p-SMR and p-HEIDI > 0.05 was retained as representative—38 high-confidence gene-probe associations were retained (Supplementary Table 2). Volcano plot visualization of 38 optic neuritis-associated loci revealed 18 significantly dysregulated genes(FDR<0.05), comprising 11 upregulated risk genes and 7 downregulated protective genes (Supplementary Table 3).Most potent risk drivers(red) (p<0.001) were HLA-DRB1 (β=1.193),KDM5B (β=1.525) and TCF25 (β=1.413),Strongest protectors(blue)(p<0.001) were CYP21A1P (β=-2.349),FLOT1 (β=-1.773) and SPG7 (β=-1.592)(Fig 1).

**Fig 1.**
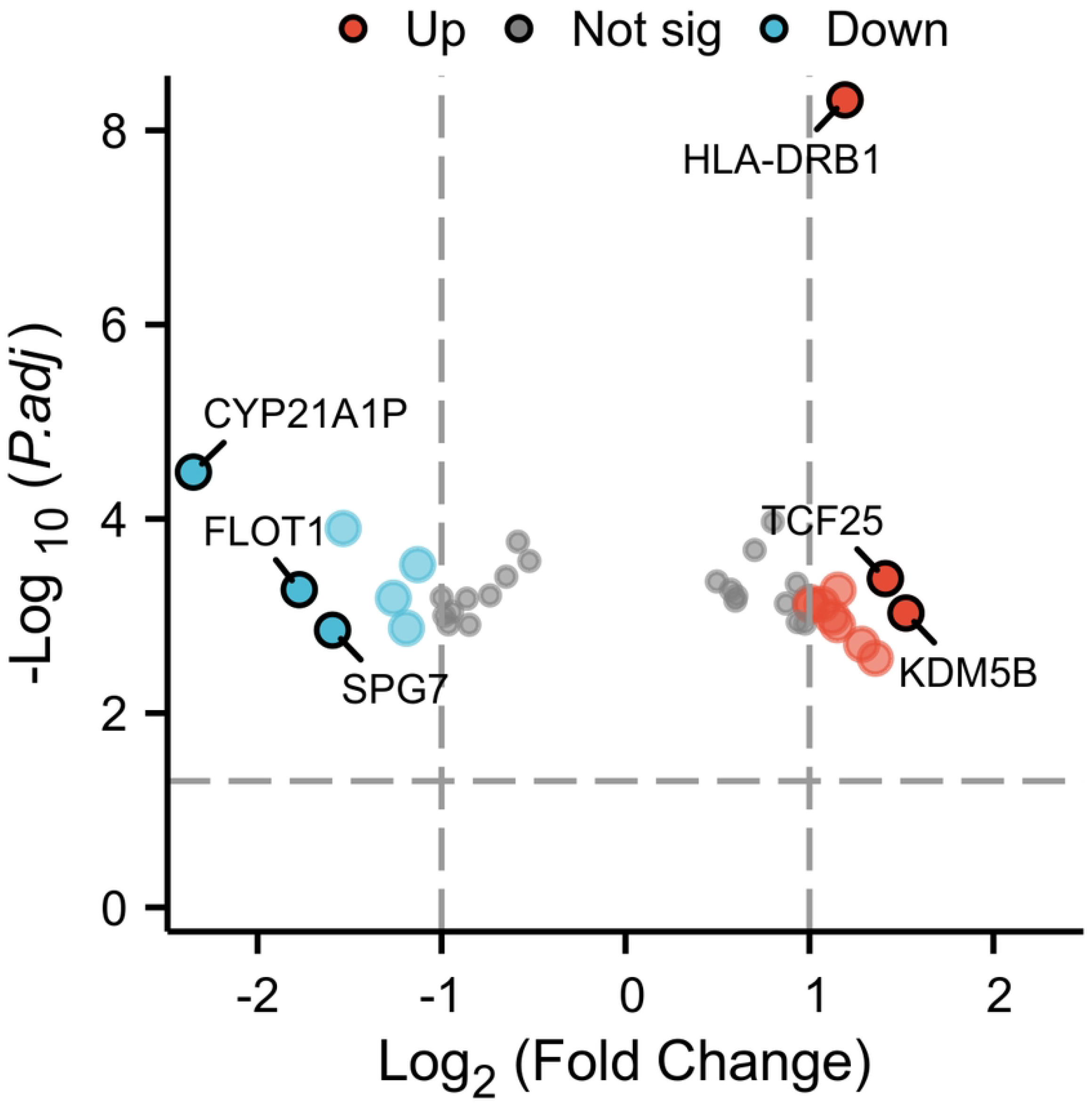
Volcano plot of optic neuritis-associated genetic variants. Significant risk/protective loci were identified from FinnGen R10 GWAS (n=38 loci, threshold p<0.001). Top risk drivers(red): HLA-DRB1(β=1.193,), KDM5B (β=1.525), TCF25_(β=1.413) Top protective variants(blud): CYP21A1P (β=-2.349), FLOT1(β=-1.773), SPG7(β=-1.592)

### 2. Functional enrichment analysis of 38 optic neuritis-associated genes (Fig 2). Key findings include(Full results: Supplementary Table 4)

#### 2.1 Gene Ontology (GO) Analysis

##### Biological Process (BP)

Significant enrichment in antigen processing/presentation of endogenous peptides (GO:0002483, p.adjust=0.0025) and regulation of T cell-mediated cytotoxicity (GO:0001916, p.adjust=0.0034). These pathways reflect CD8+ T cell activation against myelin antigens.

**Fig 2.**
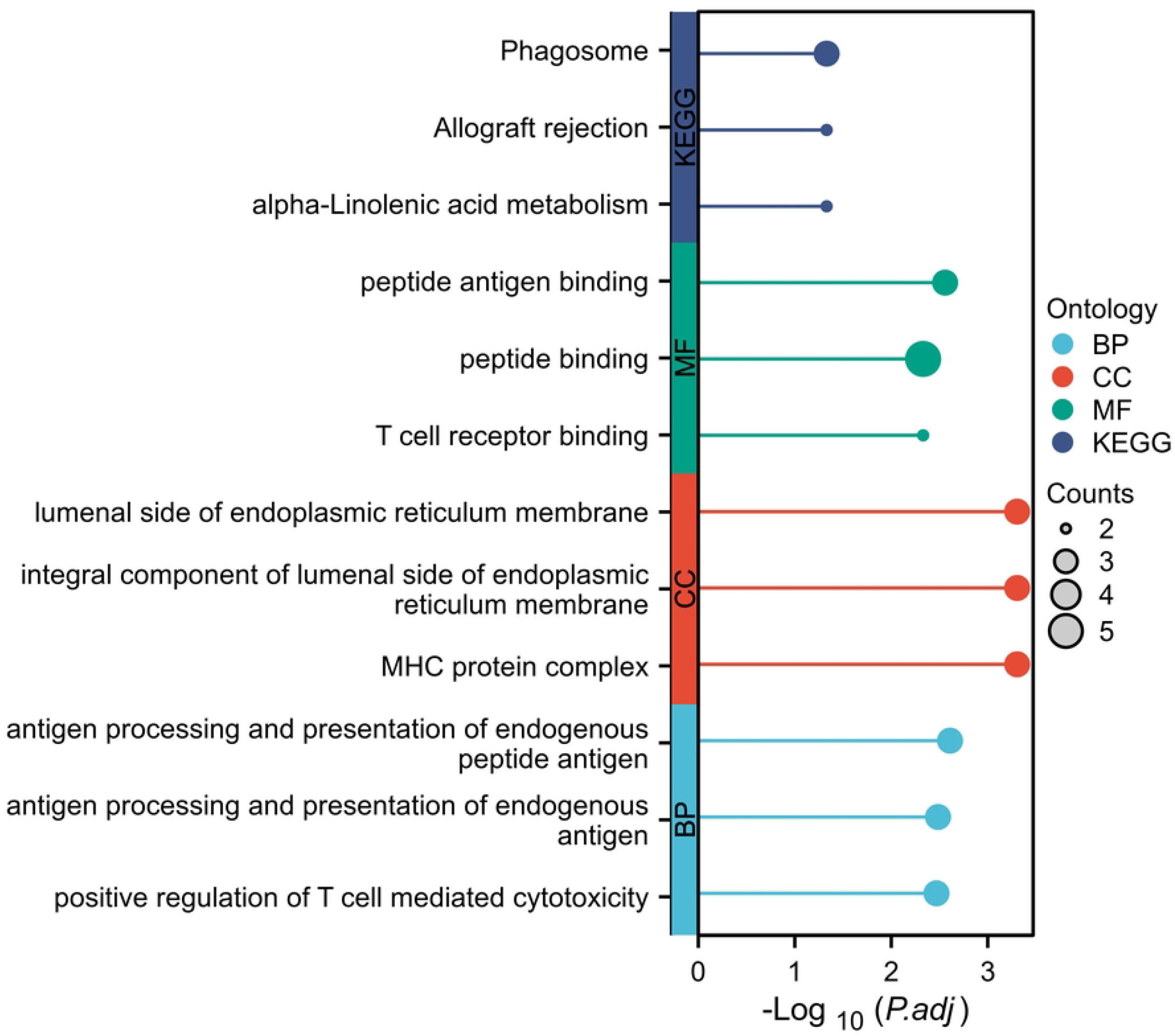
Functional enrichment analysis of 38 optic neuritis-associated genes. Significantly enriched terms (FDR < 0.05) are shown for: Molecular functions(MF): Peptide antigen binding, T cell receptor binding.Cellular components(CC): ER lumenal membrane, MHC complex, Reticulum membrane.Biological processes(BP): Antigen processing/presentation, T cell cytotoxicity regulation.KEGG pathways: Phagosome, Allograft rejection, α-Linolenic acid metabolism.

##### Cellular Component (CC)

Genes localized to the MHC protein complex (GO:0042611, p.adjust=0.0005) and endoplasmic reticulum membrane (GO:0071556, p.adjust=0.0005), consistent with astrocyte-mediated antigen presentation in ON lesions.

##### Molecular Function (MF)

Enriched in peptide antigen binding (GO:0042605, p.adjust=0.0028) and T cell receptor binding (GO:0042608, p.adjust=0.0047), implicating HLA-TCR interactions in ON pathogenesis.

Molecular Function (MF): Enriched in peptide antigen binding (GO:0042605, p.adjust=0.0028) and T cell receptor binding (GO:0042608, p.adjust=0.0047), implicating HLA-TCR interactions in ON pathogenesis.

### 2.2 KEGG Pathway Analysis

hsa05332 (Graft-versus-host disease) and hsa05165 (Human papillomavirus infection) were significantly enriched (p.adjust=0.0468), indicating shared mechanisms between autoimmune ON and transplant rejection.

hsa04145 (Phagosome) enrichment (p.adjust=0.0468) suggests microglial clearance of myelin debris amplifies inflammation, as observed in demyelinating ON models.

#### 3. Network Pharmacology for Drug Repurposing

Based on the analysis of drug-target gene interactions, a total of 1,847 potential drugs were screened (complete list in Supplementary Table 5). Among them, Omacetaxine mepesuccinate showed the most statistically significant interaction with ribosomal protein genes RPL3/rpl2 (Z-score = -4.186, p = 1.42e-5, FDR = 0.024), and this result passed multiple testing correction. The remaining candidate drugs did not reach the significance threshold due to FDR >0.05. Figure 3 illustrates the drug-target-gene interaction relationships through a network diagram.

**Fig 3.**
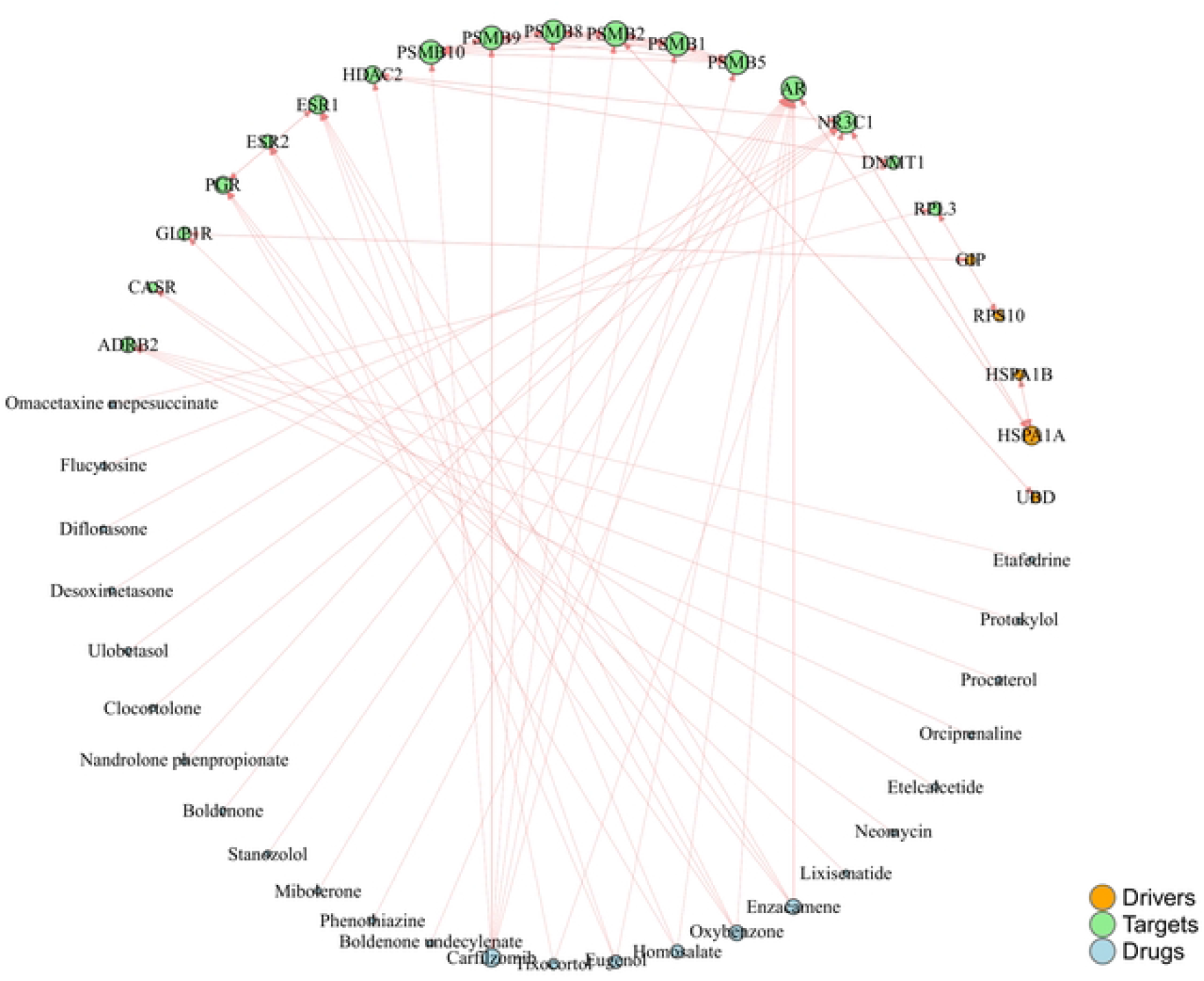
Network analysis of drug-target-gene interactions in optic neuritis. The diagram visualizes relationships between candidate drugs (e.g.Omacetaxine mepesuccinate, Flucytosine), their molecular targets (e.g.PSMB10, DNMT1, NR3C1), and disease-associated genes (e.g.ESR1, HSPA1A).

## Discussion

This study represents the first integrated analysis of GWAS, functional genomics, and network pharmacology to elucidate the genetic architecture of optic neuritis (ON) and identify repositionable therapeutics. Our findings establish three principal advances in understanding ON pathobiology:

### 1. ON Genetic Susceptibility Anchored in Antigen Presentation Machinery

The robust association of HLA-DRB1 (OR=3.297, FDR<0.01) with ON risk, together with the enrichment of MHC complex components (GO:0042611, FDR=0.0005), corroborates seminal work linking HLA haplotypes to inflammatory demyelination [9]. HLA-DRB1 facilitates myelin-reactive T cell activation [9,14], supported by intrathecal anti-myelin antibody synthesis in acute ON [15]. Notably, the Jordanian MS cohort exhibited a specific association of HLA-DRB1 alleles with ON patients [9], aligning with our identification of HLA-DRB1 as a top risk locus. The colocalization of antigen processing genes within the endoplasmic reticulum membrane (GO:0071556, FDR=0.0005) further reinforces pathological models. In these models, astrocyte ER stress amplifies autoimmune responses [16].

### 2. Convergent Pathways: From Viral Mimicry to Microglial Activation

Our pathway analysis reveals striking parallels between autoimmune ON and allogeneic rejection responses. The enrichment of graft-versus-host disease (hsa05332, FDR=0.0468) and HPV infection (hsa05165, FDR=0.0468) pathways supports the “molecular mimicry” hypothesis [17], wherein viral antigens trigger cross-reactive immune attacks on optic nerve myelin. Complementing this, activation of the phagosome pathway (hsa04145, FDR=0.0468) suggests that microglial clearance of myelin debris generates pro-inflammatory feedback loops. This mechanism has been experimentally validated in MHV-induced ON models [18].

### 3. Ribosomal-Targeted Therapy: A Mechanistically Grounded Candidate

Through stringent network pharmacology filtering, omacetaxine mepesuccinate emerged as the sole statistically supported repurposing candidate (FDR = 0.0467) targeting ribosomal proteins RPL3 and RPL2. This finding is mechanistically congruent with viral proteins subverting axonal ribosomal function in experimental optic neuritis (ON) [19].Omacetaxine mepesuccinate, or homoharringtonine (HHT), is a semi-synthetic alkaloid derived from Cephalotaxus plants. It is recognized for its role as a protein synthesis inhibitor, specifically targeting the 60S ribosomal subunit essential for cellular protein production [20]. This function makes it a promising treatment for various cancers, including chronic myeloid leukemia (CML) and acute myeloid leukemia (AML) [21], with recent studies extending its potential to acute lymphoblastic leukemia (ALL) and even solid tumors such as triple-negative breast cancer(TNBC) [22]. By inhibiting protein synthesis, Omacetaxine mepesuccinate could also be a strategic option for treating optic neuritis by reducing immune cell activation and antigen presentation, key factors in the disease’s pathology [23]. Beyond oncology, HHT’s potential in combating viral infections and immune-related diseases has garnered interest, with research indicating its effectiveness in inhibiting viral replication [24, 25]. Thus, HHT warrants further investigation for its possible role in treating optic neuritis triggered by viral antigens.

### 4. Clinical Implications and Limitations

Despite the statistical rigor of our analysis, limitations exist:The FinnGen cohort’s predominantly European ancestry may limit generalizability.Functional validation of Omacetaxine’s neuroprotective potential in ON models is warranted.Pathway analyses implicate but cannot prove causality in microglial activation.

## Conclusion

By establishing HLA-DRB1 and ribosomal dysfunction as pivot points in ON pathogenesis, this study bridges genetic risk with actionable therapeutic strategies. The prioritization of Omacetaxine – an FDA-approved agent – offers a rapid translation pathway. Future studies should validate its efficacy in representative ON models while exploring synergies with agents like dimethyl fumarate that target complementary pathways.

The completed Human Participants Research Checklist is provided as Supporting Information (S1_Checklist). All GWAS summary statistics are available in the FinnGen database (accession code: finn-b-H7_OPTNEURITIS).

## Funding

The author(s) received no specific funding for this work

## Author contributions

**Conceptualization:** Yan Zhang, Yihong Huang.

**Data curation:** Maosheng Guo, Yan Zhang.

**Investigation:** Yan Zhang, Maosheng Guo.

**Methodology:** Yan Zhang, Maosheng Guo, Yihong Huang.

**Project administration:** Yihong Huang.

**Supervision:** Yihong Huang.

**Validation:** Maosheng Guo, Yan Zhang.

**Writing – original draft:** Yan Zhang.

**Writing – review & editing:** Yan Zhang, Maosheng Guo, Yihong Huang.

## Notes

### Competing Interest Statement

The authors have declared no competing interest.

## References

[1] Chuenkongkaew W, Chirapapaisan N. Optic neuritis: characteristics and visual outcome. J Med Assoc Thai. 2003 Mar;86(3):238-43. PMID: 12757063.

[2] Duvigneaud Z, Lardeux P, Verrecchia S, Benyahya L, Marignier R, Froment Tilikete C. Diagnostic criteria for optic neuritis in the acute and subacute phase: clinical uses and limitations. J Neurol. 2024 Aug;271(8):5629–5636. doi: 10.1007/s00415-024-12540-9. Epub 2024 Jul 2. PMID: 38954036.

[3] Pula JH, Macdonald CJ. Current options for the treatment of optic neuritis. Clin Ophthalmol. 2012;6:1211–23. doi: 10.2147/OPTH.S28112. Epub 2012 Jul 31. PMID: 22927730; PMCID: PMC3422147.

[4] Petzold A, Fraser CL, Abegg M, Alroughani R, Alshowaeir D, Alvarenga R, et al. Diagnosis and classification of optic neuritis. Lancet Neurol. 2022 Dec;21(12):1120–1134. doi: 10.1016/S1474-4422(22)00200-9. Epub 2022 Sep 27. PMID: 36179757.

[5] Fernández Blanco L, Marzin M, Leistra A, van der Valk P, Nutma E, Amor S. Immunopathology of the optic nerve in multiple sclerosis. Clin Exp Immunol. 2022 Aug 19;209(2):236–246. doi: 10.1093/cei/uxac063. PMID: 35778909; PMCID: PMC9390848.

[6] Xu LT, Bermel RA, Nowacki AS, Kaiser PK. Optical Coherence Tomography for the Detection of Remote Optic Neuritis in Multiple Sclerosis. J Neuroimaging. 2016 May;26(3):283–8. doi: 10.1111/jon.12326. Epub 2016 Jan 15. PMID: 26773711.

[7] Naphattalung Y, Chuenkongkaew WL, Chirapapaisan N, Laowanapiban P, Sawangkul S. Plasma exchange for acute optic neuritis in neuromyelitis optica or neuromyelitis optica spectrum disorder: a systematic review. Ann Med. 2023 Dec;55(1):2227422. doi: 10.1080/07853890.2023.2227422. PMID: 37387119; PMCID: PMC10316725.

[8] Qiu X, Huang MN, Ping S. Genetic susceptibility and causal pathway analysis of eye disorders coexisting in multiple sclerosis. Front Immunol. 2024 Feb 5;15:1337528. doi: 10.3389/fimmu.2024.1337528. PMID: 38375484; PMCID: PMC10875133.

[9] Khdair SI, Al-Khareisha L, Abusara OH, Hammad AM, Khudair A. HLA-class II genes association with multiple sclerosis: An immunogenetic prediction among multiple sclerosis Jordanian patients. PLoS One. 2025 Feb 25;20(2):e0318824. doi: 10.1371/journal.pone.0318824. PMID: 39999097; PMCID: PMC11856260.

[10] Duvigneaud Z, Lardeux P, Verrecchia S, Benyahya L, Marignier R, Froment Tilikete C. Diagnostic criteria for optic neuritis in the acute and subacute phase: clinical uses and limitations. J Neurol. 2024 Aug;271(8):5629–5636. doi: 10.1007/s00415-024-12540-9. Epub 2024 Jul 2. PMID: 38954036.

[11] França LC, Fontes-Dantas FL, Garcia DG, de Araújo AD, da Costa Gonçalves JP, et al. Molecular mimicry between Zika virus and central nervous system inflammatory demyelinating disorders: the role of NS5 Zika virus epitope and PLP autoantigens. Arq Neuropsiquiatr. 2023 Apr;81(4):357–368. doi: 10.1055/s-0043-1768698. Epub 2023 May 9. PMID: 37160141; PMCID: PMC10169219.

[12] Wu T, Jiang H, Lin C, Peng J, Kong X, Yu J, Wang J, Cui S. Gut microbial profiles of patients with optic neuritis or myasthenia gravis. J Int Med Res. 2025 Feb;53(2):3000605251314817. doi: 10.1177/03000605251314817. PMID: 39904582; PMCID: PMC11795606.

[13] Guney E, Menche J, Vidal M, Barábasi AL. Network-based in silico drug efficacy screening. Nat Commun. 2016 Feb 1;7:10331. doi: 10.1038/ncomms10331. PMID: 26831545; PMCID: PMC4740350.

[14] Lai CT, Li W, Sun YB, Meng C, Chi KM, Lu CF, Yu HF, Zhang ZX. [Clinical characteristic of optic neuritis and its relevancy with human leukocyte antigen]. Zhonghua Yan Ke Za Zhi. 2006 Jun;42(6):501-6. Chinese. PMID: 16857128.

[15] Sellebjerg F, Madsen HO, Frederiksen JL, Ryder LP, Svejgaard A. Acute optic neuritis: myelin basic protein and proteolipid protein antibodies, affinity, and the HLA system. Ann Neurol. 1995 Dec;38(6):943–50. doi: 10.1002/ana.410380616. PMID: 8526468.

[16] Chun BY, Kim JH, Nam Y, Huh MI, Han S, Suk K. Pathological Involvement of Astrocyte-Derived Lipocalin-2 in the Demyelinating Optic Neuritis. Invest Ophthalmol Vis Sci. 2015 Jun;56(6):3691–8. doi: 10.1167/iovs.15-16851. PMID: 26047170.

[17] Tselis AC, Lisak RP. Acute disseminated encephalomyelitis and isolated central nervous system demyelinative syndromes. Curr Opin Neurol. 1995 Jun;8(3):227–9. doi: 10.1097/00019052-199506000-00012. PMID: 7551123.

[18] Shindler KS, Chatterjee D, Biswas K, Goyal A, Dutt M, Nassrallah M, Khan RS, Das Sarma J. Macrophage-mediated optic neuritis induced by retrograde axonal transport of spike gene recombinant mouse hepatitis virus. J Neuropathol Exp Neurol. 2011 Jun;70(6):470–80. doi: 10.1097/NEN.0b013e31821da499. PMID: 21572336; PMCID: PMC3110774.

[19] Zyla K, Larabee CM, Georgescu C, Berkley C, Reyna T, Plafker SM. Dimethyl fumarate mitigates optic neuritis. Mol Vis. 2019 Aug 22;25:446-461. PMID: 31523122; PMCID: PMC6707756.

[20] Völkl M, Burgers LD, Zech TJ, Ciurus S, Dorovska S, Liu H, Zahler S, Fürst R. Homoharringtonine (omacetaxine mepesuccinate) limits the angiogenic capacity of endothelial cells and reorganises filamentous actin. Biomed Pharmacother. 2025 May;186:118025. doi: 10.1016/j.biopha.2025.118025. Epub 2025 Apr 4. PMID: 40184838.

[21] Quintás-Cardama A, Cortes J. Omacetaxine mepesuccinate--a semisynthetic formulation of the natural antitumoral alkaloid homoharringtonine, for chronic myelocytic leukemia and other myeloid malignancies. IDrugs. 2008 May;11(5):356-72. PMID: 18465678.

[22] Plett R, Mellor P, Kendall S, Hammond SA, Boulet A, Plaza K, Vizeacoumar FS, Vizeacoumar FJ, Anderson DH. Homoharringtonine demonstrates a cytotoxic effect against triple-negative breast cancer cell lines and acts synergistically with paclitaxel. Sci Rep. 2022 Sep 19;12(1):15663. doi: 10.1038/s41598-022-19621-7. PMID: 36123435; PMCID: PMC9485251.

[23] Chen J, Hui Y, Zhai Y, Yang M, Zhang X, Mi Y, Wang J, Wei H. Serum albumin is associated with the inherent property of acute myeloid leukemia and correlates with patient outcomes. Blood Sci. 2024 May 10;6(2):e00189. doi: 10.1097/BS9.0000000000000189. PMID: 38742239; PMCID: PMC11090624.

[24] Cao J, Lu G, Wen L, Luo P, Huang Y, Liang R, Tang K, Qin Z, Chan CC, Chik KK, D. J, Yin F, Ye ZW, Chu H, Jin DY, Yuen KY, Chan JF, Yuan S. Severe fever with thrombocytopenia syndrome virus (SFTSV)-host interactome screen identifies viral nucleoprotein-associated host factors as potential antiviral targets. Comput Struct Biotechnol J. 2021 Oct 1;19:5568–5577. doi: 10.1016/j.csbj.2021.09.034. PMID: 34712400; PMCID: PMC8523828.

[25] Zheng X, Chen J, Zhang Y, Hu S, Bi C, Singla RK, Kamal MA, Horimoto K, Shen B. Translational Informatics Driven Drug Repositioning for Neurodegenerative Disease. Curr Neuropharmacol. 2025 Feb 6. doi: 10.2174/011570159X327908241121062335. Epub ahead of print. PMID: 39936420.

